# Nijmegen Breakage Syndrome (NBS) is a Telomeropathy: Analysis of Telomere Length in NBS Homo- and Heterozygotes and Humanized Nbs Mice

**DOI:** 10.1101/571026

**Authors:** Raneem Habib, Ryong Kim, Heidemarie Neitzel, Ilja Demuth, Krystyna Chrzanowska, Eva Seemanova, Renaldo Faber, Martin Digweed, Kathrin Jäger, Karl Sperling, Michael Walter

## Abstract

The autosomal recessive genetic disorder Nijmegen breakage syndrome (NBS) is characterized by a defect in DNA double-strand break repair protein nibrin and chromosome instability associated with a high predisposition to cancer. Here we hypothesized that impaired nibrin/MRE11/RAD50 telomere maintenance complex may also affect telomere length and modulate the cancer phenotype.

Telomere length was studied in blood from 38 homozygous and 27 heterozygous individuals, in one homozygous fetus, and in sex NBS lymphoblastoid cell lines (all with the founder mutation c.657_661del5), and in three humanized Nbs mice, using qPCR, TRF and Q-FISH.

Telomere lengths were markedly but uniformly reduced to 20-40% of healthy controls. There was no correlation between telomere length and severity of clinical phenotype or age of death. By contrast, individual patients with very short telomeres displayed long survival times after cancer manifestation. Mildly accelerated telomere attrition was found in older NBS heterozygotes. In the NBS-fetus, the spinal cord, brain and heart had the longest telomeres, skin the shortest. Humanized Nbs mice (with much longer telomeres than those in human beings) did not show accelerated telomere attrition.

Our data clearly show that NBS is a secondary telomeropathy with unique features. Telomere attrition in NBS may cause genetic instability and contribute to the high cancer incidence in NBS. On the other hand, short telomeres may prevent an even worse phenotype when a tumor has developed. These data may help to understand the high cancer rate in NBS and also the bifunctional role of telomere shortening in cancerogenesis.

**Author Summary:** DNA damage is harmful because it leads to mutations in genes that initiate or accelerate cancerogenesis. The devastating consequences of DNA damage are manifested in diseases with non-functional repair pathways such as Nijmegen breakage syndrome (NBS). A common feature of these diseases is a high tumor incidence. However, cancer incidence varies and is not clear why it is highest for NBS. In a previous study, we have shown that the underlying nebrin mutation not only leads to defective DNA repair but also to higher degree of oxidative stress that generates further DNA lesions. Nibrin may play also an important role in protecting chromosome ends, the telomeres, from inappropriate DNA repair. Therefore we examined the telomere length in NBS and show markedly reduced values in affected patients but not in NBC mice (with much milder phenotype and longer telomeres). Telomere attrition contributes to genetic instability and may thus contribute to the high cancer incidence in NBS. Individual patients with very short telomeres, however, displayed long survival times after cancer manifestation. Thus, short telomeres may also prevent an even worse phenotype when a tumor has developed. These data are fundamental to understanding the high cancer rate in NBS and also the bifunctional role of telomere shortening in cancer.

## Introduction

The autosomal recessive genetic disorder Nijmegen breakage syndrome (NBS) was first described in 1981 in two patients living in Nijmegen, Holland [1]. It is characterized by chromosome instability associated with microcephaly, immunodeficiency, hypersensitivity to X-irradiation, and a high predisposition to cancer [2-4]. Our group mapped the gene for NBS to chromosome 8q21 in 1997 [5] and identified nibrin, a DNA double-strand break repair protein, as defective protein one year later [6]. More than 90% of NBS patients are homozygous for a founder mutation, c.657_661del5, in the NBN gene [6-9]. By the age of 20 years, more than 40% of patients have developed a malignant disease, predominantly of lymphoid origin. Even heterozygous carriers of the founder mutation have an increased cancer risk [9,10].

Nibrin, the product of the *NBN(NBS1)* gene, is part of the MRE11/RAD50 (MRN) complex that is involved in the repair of DNA double strand breaks (DSBs), both by homologous recombination repair (HRR) and non-homologous end-joining (NHEJ), the processing of DSBs in immune gene rearrangements, and meiotic recombination[11-14]. The important role of this complex in mediating the ATM-dependent repair of DSBs likely explains the high predisposition to cancer in NBS.

However, cancer incidence varies among genetic instability syndromes and is not clear why it is highest in NBS. We have recently shown that truncated nibrin protein in NBS leads to high levels of reactive oxygen after DNA damage and that this increased oxidative stress is caused by depletion of NAD^+^ due to hyperactivation of the strand-break sensor, Poly(ADP-ribose) polymerase. Data suggested that the extremely high incidence of malignancy among NBS patients is the result of the combination of a primary DSB repair deficiency with secondary changes such as oxidative DNA damage [15].

Nibrin is multifunctional and may play also an important role in protecting chromosome ends, the telomeres, from inappropriate DNA repair. Telomeric DNA is an evolutionary highly conserved repetitive sequence which plays a crucial role both in cellular senescence and in carcinogenesis. Nibrin interacts with TRF1 and is part of the shelterin complex regulating telomere length. In addition, it is associated with TRF2, which binds to the telomere and participates in formation and stabilization of T-loops, which are required for telomere replication [16,17]. Moreover, nibrin was found to be involved in telomerase-independent telomere maintenance pathway, the so-called alternative lengthening of telomeres (ALT) mechanism[18].

We therefore analysed telomere length in blood cells and lymphoblastoid cell lines from NBS patients homozygous for the common founder mutation, in heterozygote individuals, in an NBS fetus, and in a humanized Nbs mouse which carries the human NBN gene with the founder mutation. Based on the described experimental findings and previously reported shorter telomeres in individual cases and accelerated senescence of NBS fibroblasts in culture [19-23] we hypothesized that NBS is a secondary telomeropathy or even belongs to an impaired telomere maintenance spectrum disorder (ITM), according to the recently proposed nomenclature [24], and that telomere abnormalities may accelerate cancer manifestation. A number of similar genetic disorders may also display dysfunctional telomeres, a defective DNA damage response, limited cell proliferation capacity *in vitro* and an increased cancer risk [24-26].

Three different complementary methods were applied to measure telomere length: qPCR, to estimate the relative total length of telomeric repeats, Q-FISH, to determine the relative length of individual telomeres, and TRF (Terminal Restriction Fragment) analysis, to estimate the total absolute length of telomeric DNA.

## Results

### Estimation of telomere length by qPCR

Relative leukocyte telomere lengths (rLTL) of blood DNA from 38 NBS homozygotes, 27 heterozygotes, and 108 control individuals were measured by qPCR (Supplemental Tables 1-3). The regression curves show a similar decline for all three cohorts and an inverse relationship between telomere length and age (Fig. 2). When standardized for age, the curve of the NBS homozygotes shows the most distinct decline (Fig. 2, bottom).

**Fig. 1:**
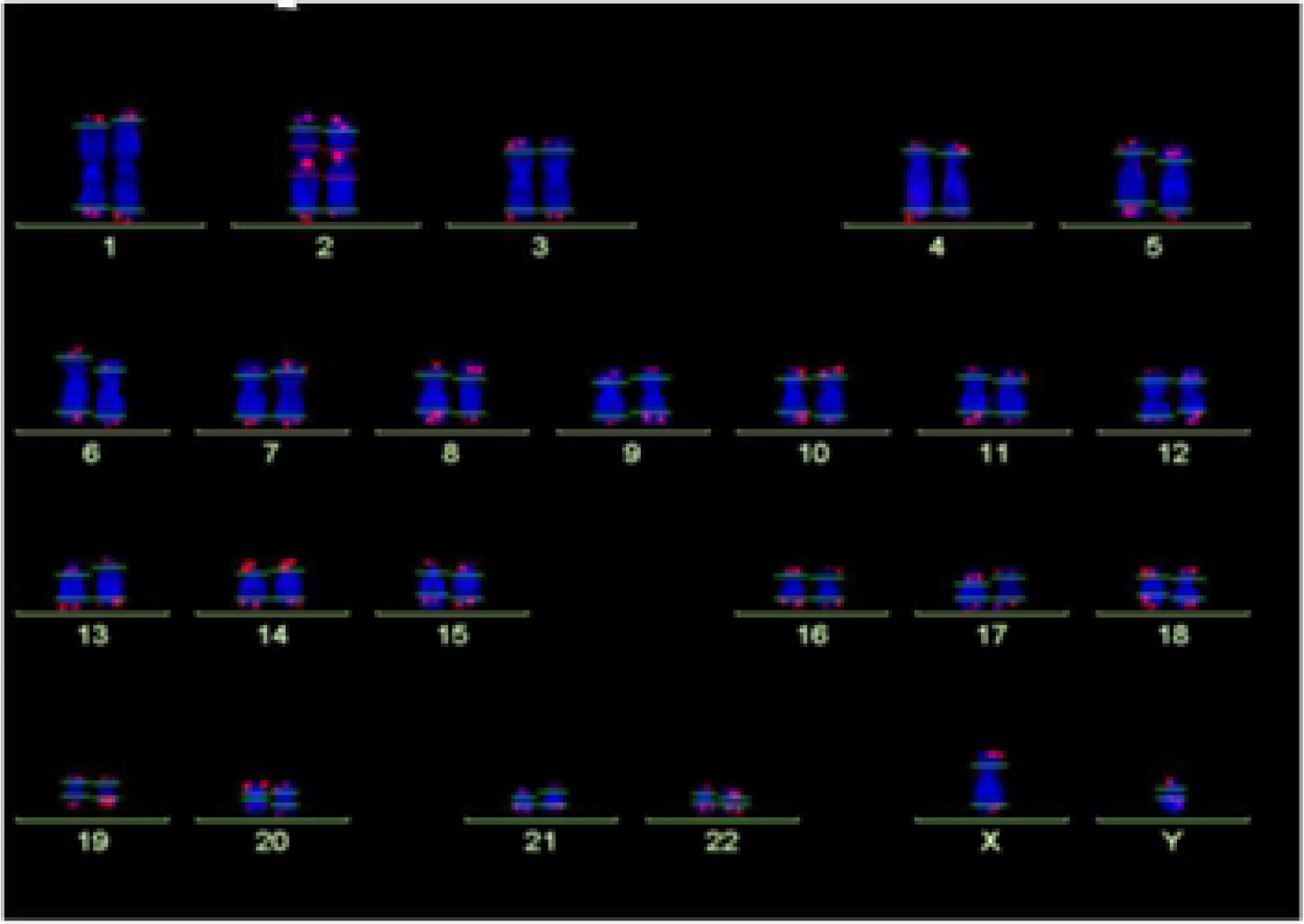
Analysis of telomere length by Q-FISH. Normal karyogramme with single telomeres stained for telomere repeats (Q-FISH) and a repetitive region in the centromeric region of chromosome 2. The horizontal lines overlaid on each chromosome define the measurement areas. Original from Habib 2012 [31].

**Fig 2:**
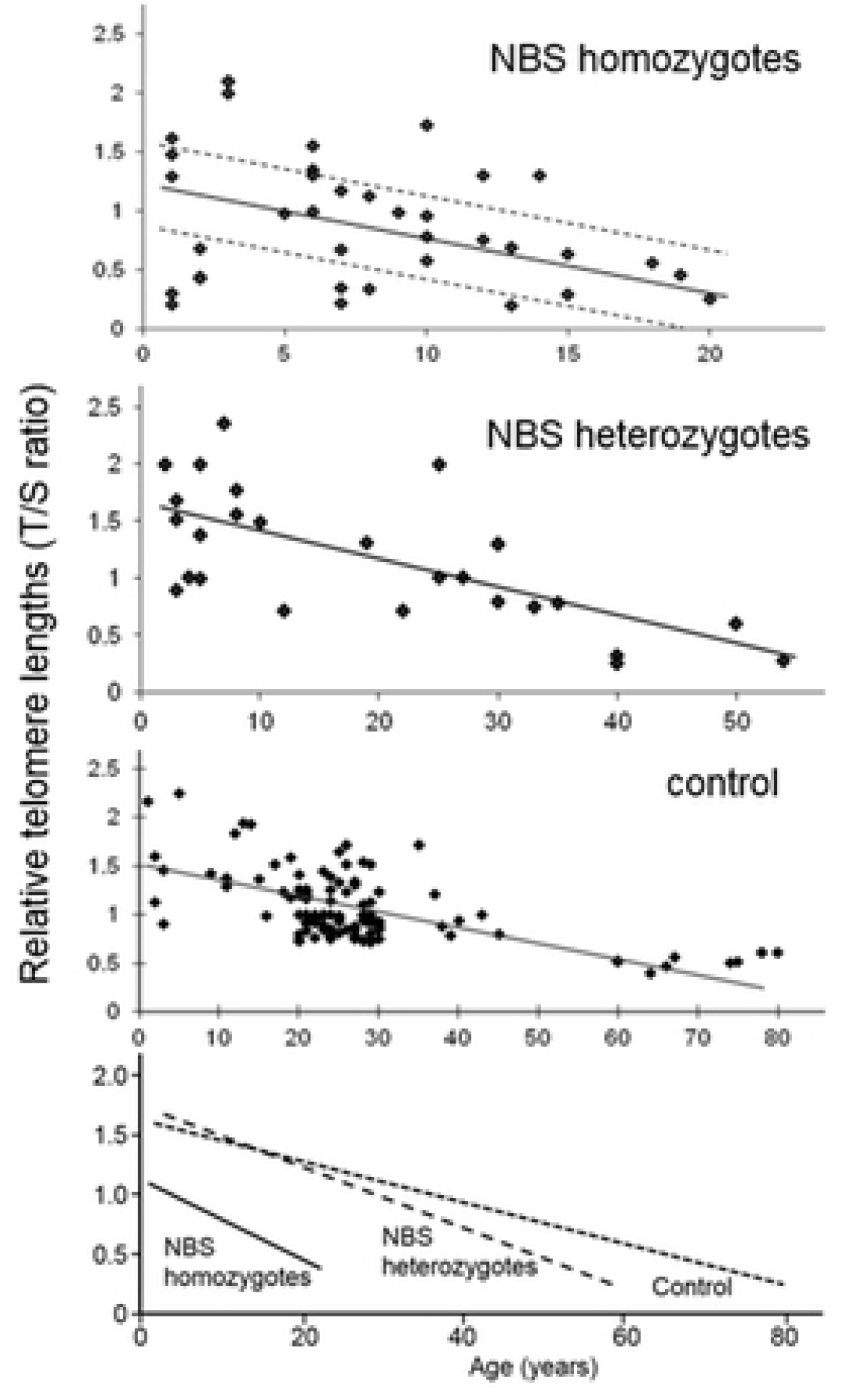
Relative telomere length as a function of age of NBS homozygotes, heterozygotes and control individuals. Relative telomere length (T/S ratio) as a function of age analysed from blood samples of 38 NBS homozygotes, 27 NBS heterozygotes and 108 control individuals by quantitative polymerase chain reaction (qPCR). The dashed lines, simply drawn by hand, separate the NBS homozygotes in those with long, medium, and short telomere length. Below: Regression curves standardized for age. Original after Habib, 2012[31].

The corresponding correlation coefficients were almost the same for controls and NBS heterozygotes, r = - 0.59 and −0.66, but much lower in NBS homozygotes with r = −0.27, which is due to the much greater variability of individual telomere lengths.

The mean relative telomere length of NBS-homozygotes was significantly shorter than that of the control individuals (P < 0.05), based on the comparison of two age matched cohorts (1 to 10, and 11 to 20 years). No significant difference was found between NBS heterozygotes and controls (Fig. 3, Supplemental Table 4).

**Fig. 3:**
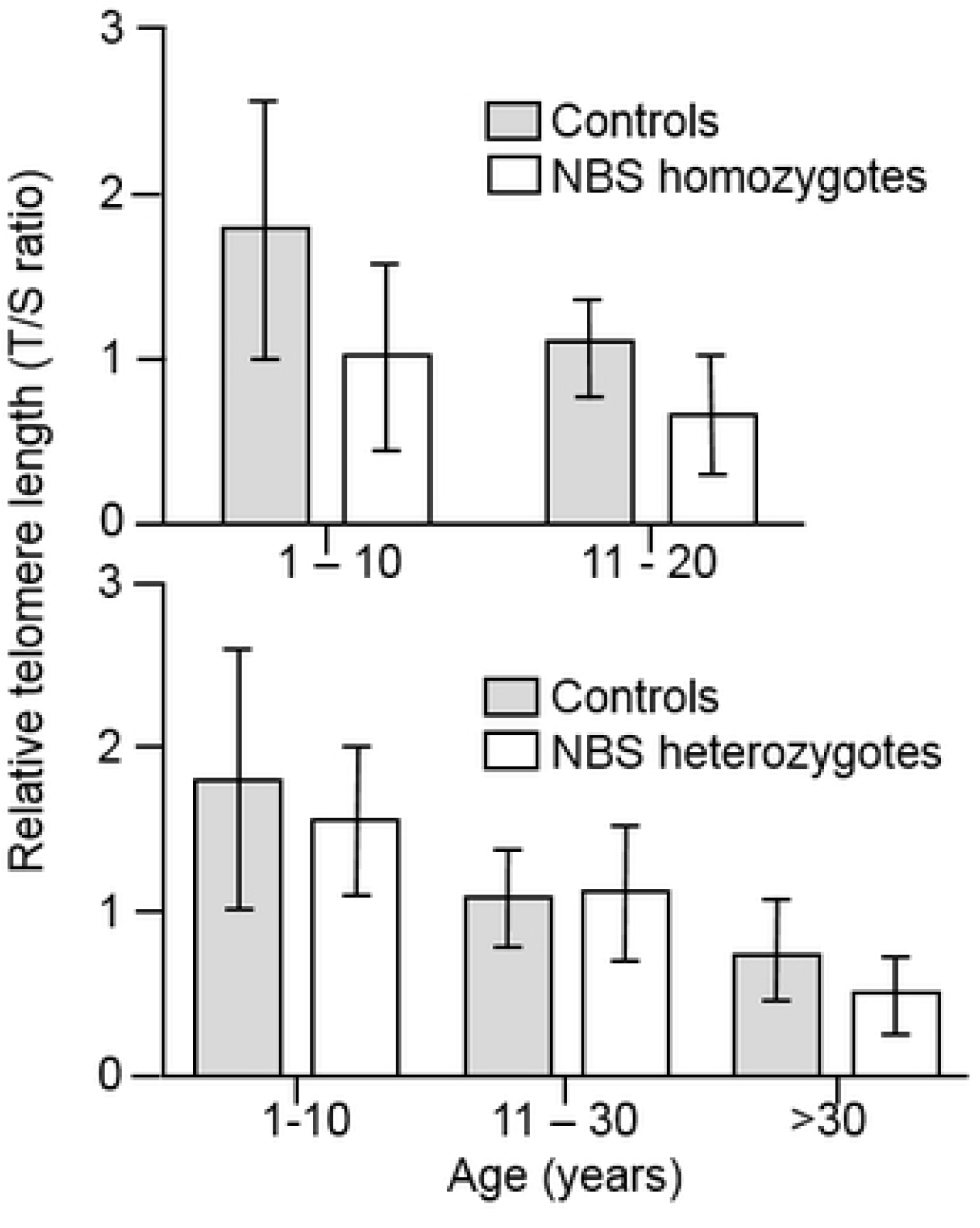
Comparison of telomere length analysed by qPCR between NBS homozygotes, heterozygotes and controls. The comparison was made for age-matched cohorts (mean values and standard deviation).

For 21 NBS homozygotes the telomere length could be correlated with clinical data, such as age at cancer manifestation and age at death. Based on Fig. 2 the individuals were classified as those with long (n=5), medium (n=8), and short (n=8) telomere length (Table 1). 17 of them had developed cancer. There was no statistical significant correlation (unpaired T-test) between telomere length and either the age at cancer manifestation or age at death (Supplemental Table 5).

**Table 1:**
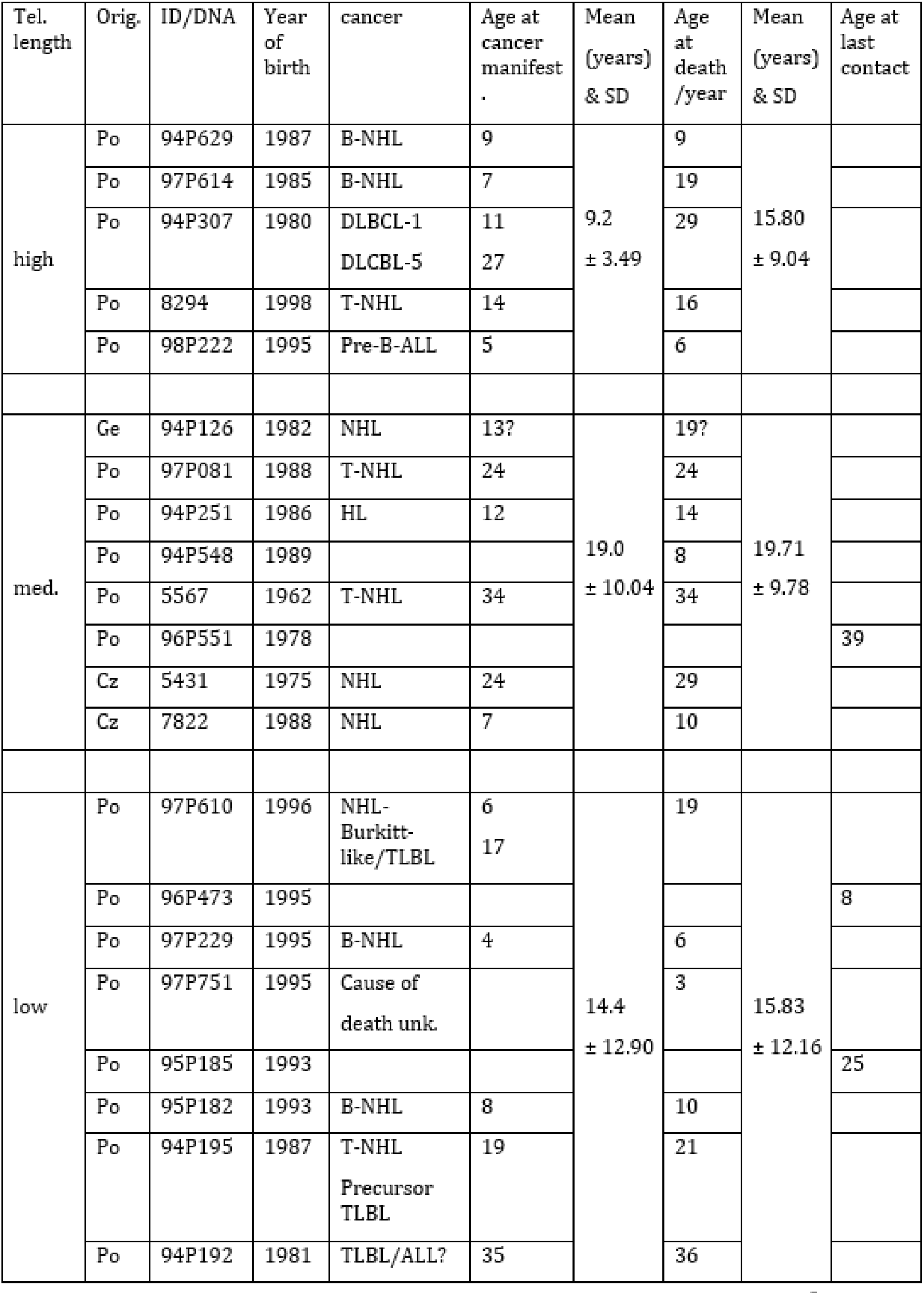
Correlation of telomere length of NBS homozygotes as classified in Fig. 2 with age at cancer manifestation and age at death. long

| Tel. length | Orig. | ID/DNA | Year of birth | cancer | Age at cancer manifest | Mean (years) & SD | Age at death /year | Mean (years) & SD | Age at last contact |
| --- | --- | --- | --- | --- | --- | --- | --- | --- | --- |
| high | Po | 94P629 | 1987 | B-NHL | 9 | 9.2<br>± 3.49 | 9 | 15.80<br>± 9.04 |  |
|  | Po | 97P614 | 1985 | B-NHL | 7 |  | 19 |  |  |
|  | Po | 94P307 | 1980 | DLBCL-1<br>DLCBL-5 | 11<br>27 |  | 29 |  |  |
|  | Po | 8294 | 1998 | T-NHL | 14 |  | 16 |  |  |
|  | Po | 98P222 | 1995 | Pre-B-ALL | 5 |  | 6 |  |  |
| med. | Ge | 94P126 | 1982 | NHL | 13? | 19.0<br>± 10.04 | 19? | 19.71<br>± 9.78 |  |
|  | Po | 97P081 | 1988 | T-NHL | 24 |  | 24 |  |  |
|  | Po | 94P251 | 1986 | HL | 12 |  | 14 |  |  |
|  | Po | 94P548 | 1989 |  |  |  | 8 |  |  |
|  | Po | 5567 | 1962 | T-NHL | 34 |  | 34 |  |  |
|  | Po | 96P551 | 1978 |  |  |  |  |  | 39 |
|  | Cz | 5431 | 1975 | NHL | 24 |  | 29 |  |  |
|  | Cz | 7822 | 1988 | NHL | 7 |  | 10 |  |  |
| low | Po | 97P610 | 1996 | NHL-Burkitt-<br>like/TLBL | 6<br>17 | 14.4<br>± 12.90 | 19 | 15.83<br>± 12.16 |  |
|  | Po | 96P473 | 1995 |  |  |  |  |  | 8 |
|  | Po | 97P229 | 1995 | B-NHL | 4 |  | 6 |  |  |
|  | Po | 97P751 | 1995 | Cause of<br>death unk. |  |  | 3 |  |  |
|  | Po | 95P185 | 1993 |  |  |  |  |  | 25 |
|  | Po | 95P182 | 1993 | B-NHL | 8 |  | 10 |  |  |
|  | Po | 94P195 | 1987 | T-NHL<br>Precursor<br>TLBL | 19 |  | 21 |  |  |
|  | Po | 94P192 | 1981 | TLBL/ALL? | 35 |  | 36 |  |  |

In addition, telomere length was measured by qPCR in 14 different tissues of a NBS-fetus, terminated at the 32 week of pregnancy (Fig. 4). The longest relative telomere length was found in spinal cord followed by brain, while the shortest length was observed in skin.

**Fig 4:**
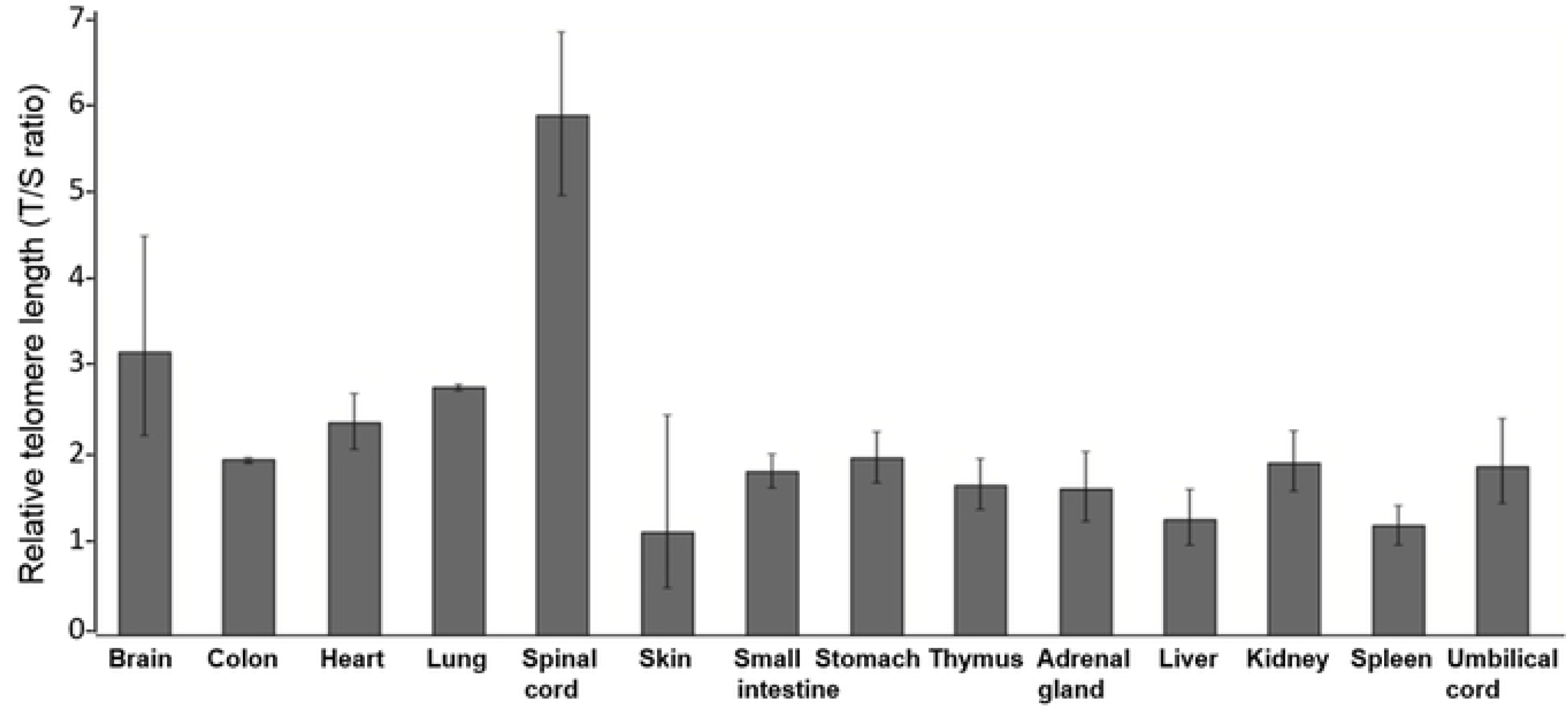
Relative telomere length of an NBS fetus. Relative telomere length ±SD (T/S ratio) in 14 different tissues of a 32 week old NBS-fetus estimated by qPCR. Original from Habib, 2012[31].

### Estimation of telomere length by Q-FISH

Telomere length was also analysed by quantitative fluorescence *in situ* hybridization of telomere repeats (Q-FISH). Based on this approach, six NBS lymphoblastoid cell lines derived from three individuals with extremely short survival after cancer manifestation (<3 years), and three with remarkably long survival (>12 years) were analysed (Table 2).

**Table 2:** Clinical data of NBS patients, donors of lymphoblastoid cell lines

| ID number/<br>sex | Cancer | Age at<br>manifest. | Age at death | Survival (years)<br>after manifestation |
| --- | --- | --- | --- | --- |
| 89P0319 ♀ | B-NHL | 7 | 7 | 0 |
| 96P0616 ♀ | B-NHL | 8.1 | 8.3 | 0.2 |
| 95P0182 ♀ | B-NHL | 8 | 10.8 | 2.8 |
| 97P0614 ♂ | B-NHL, T-ALL | 7 | 19.5 | >12 |
| RoZd ♀ | Meningeoma | 11 | >25 | >12 |
| 94P0307 ♂ | B-NHL | 11 | >25 | >12 |

The total LCL telomere length of the three patients with shorter survival after cancer manifestation was longer than of those with longer survival (Fig. 5). However, a statistically significant difference in telomere length was only found between the controls and each of the six NBS-LCLs (P<0.05; Mann-Whitney test).

**Fig 5:**
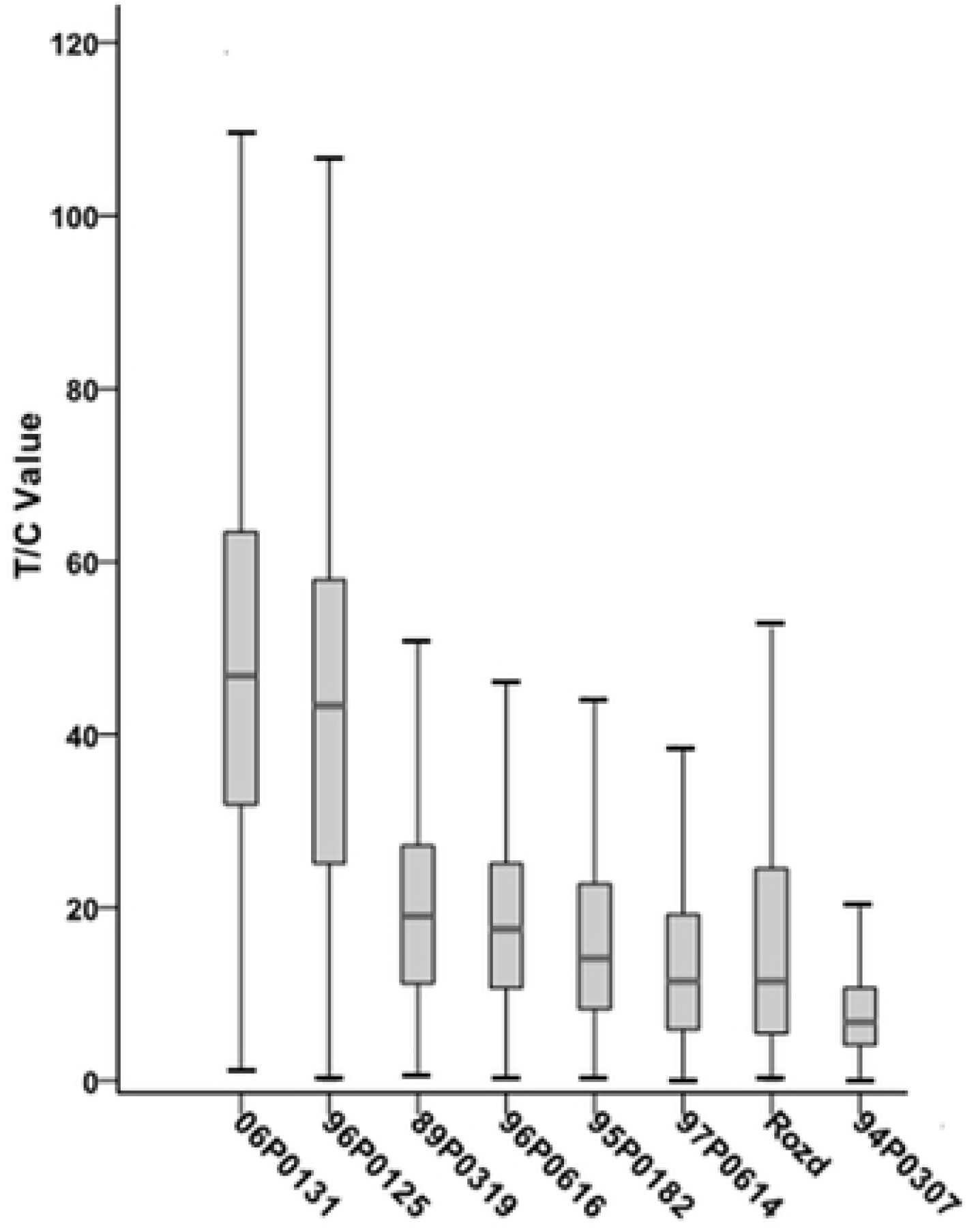
Total telomere lengths analysed by Q-FISH. Telomere length of six NBS lymphoblastoid cell lines and two control lines 06P0131 and 96P0125 analysed by Q-FISH. The boxplot presents the median, the minimum and maximum T/C values. Original from Habib, 2012[31].

The NBS cell lines exhibited some spontaneous aberrations, such as chromatid breaks and translocations. Telomere fusions were found in the cell line 94P0307 with the shortest telomeres. All individual telomeres of this line were shorter than those of the control line 06P0131 with one exception: The telomere of the p arm of one chromosome 19, which showed an enormous variability in length (Fig. 6). This was due to cellular mosaicism. In about 30% of metaphases this telomere displayed only weak fluorescence, similar to the other telomeres, however, in 70 % of the metaphases it was brightly fluorescent (Fig. 7). The difference in relative T/C values was approximately 11: 1 for the two chromosomes 19 (Table 3).

**Table 3:**
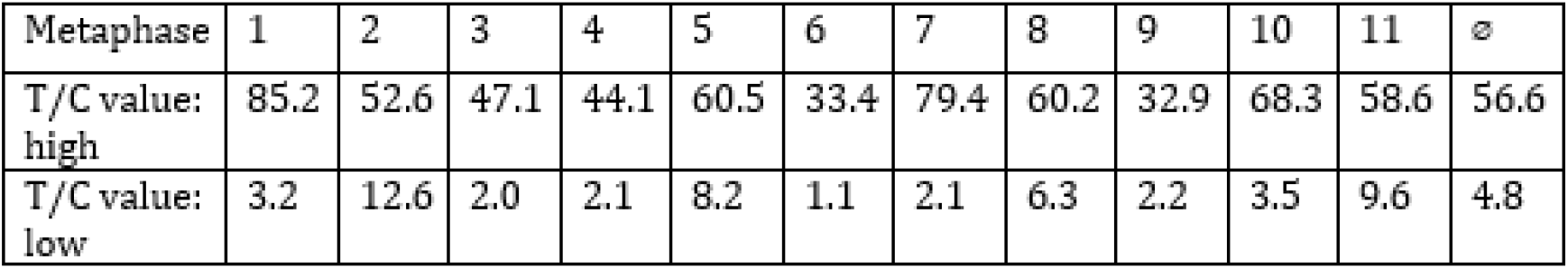
Q-FISH analysis of telomere lengths of the p arm of chromosome 19 in 11 metaphases of the NBS lymphoblastoid cell line 94P0307. T/V values are presented for the highly and weakly fluorescent chromosome 19.

**Fig 6:**
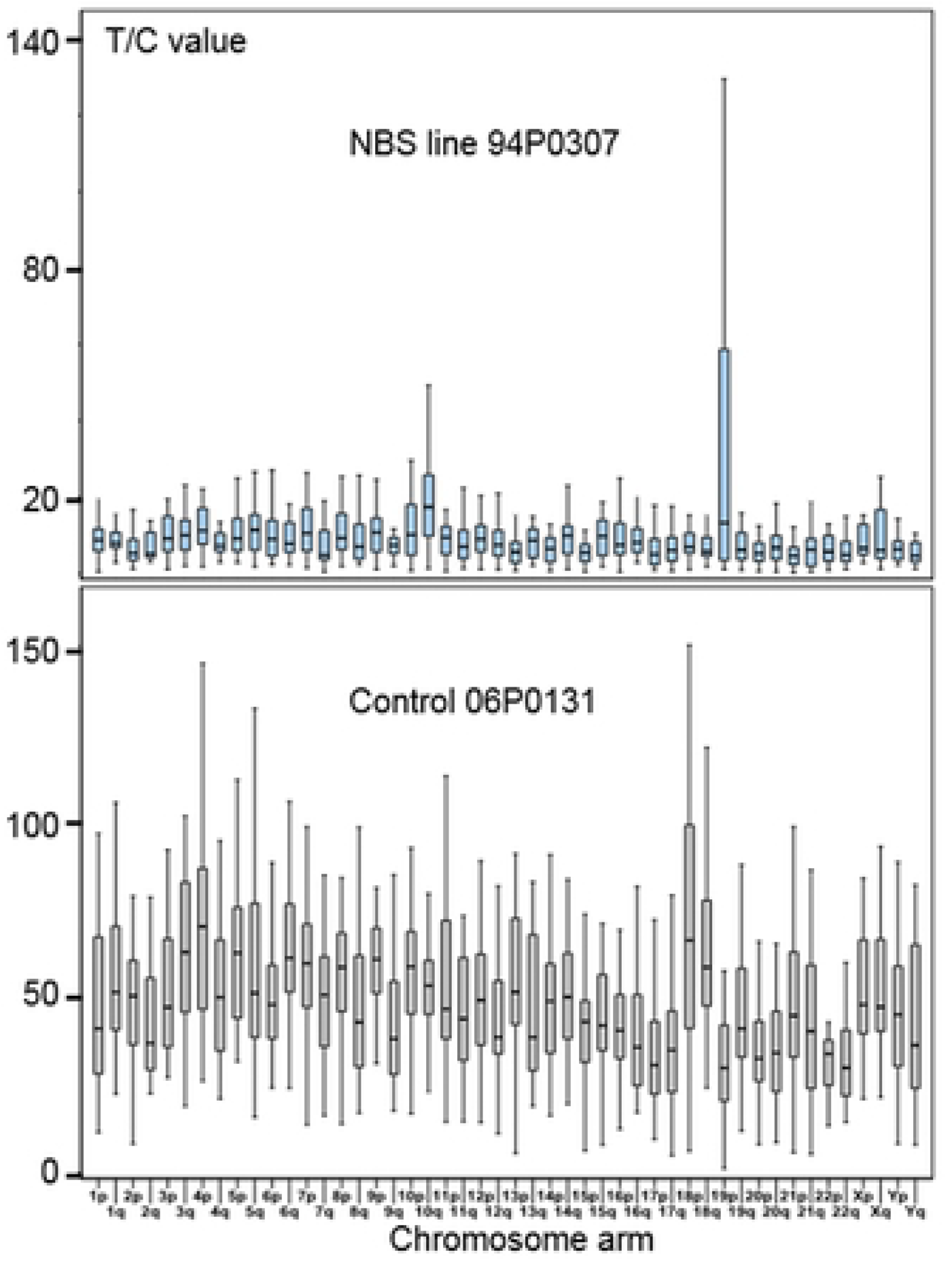
Individual telomere lengths of the NBS line 94P0307 and the control line 06P0131. Individual telomere length analysed by Q-FISH of 15 metaphases of the lymphoblastoid NBS cell line 94P0307 and a lymphoblastoid control line. The boxplot presents the median and the minimum and maximum T/C values. Note, the huge variability in telomere length of the short arm of one chromosome 19 (19p). Original from Habib, 2012[31].

**Fig. 7:**
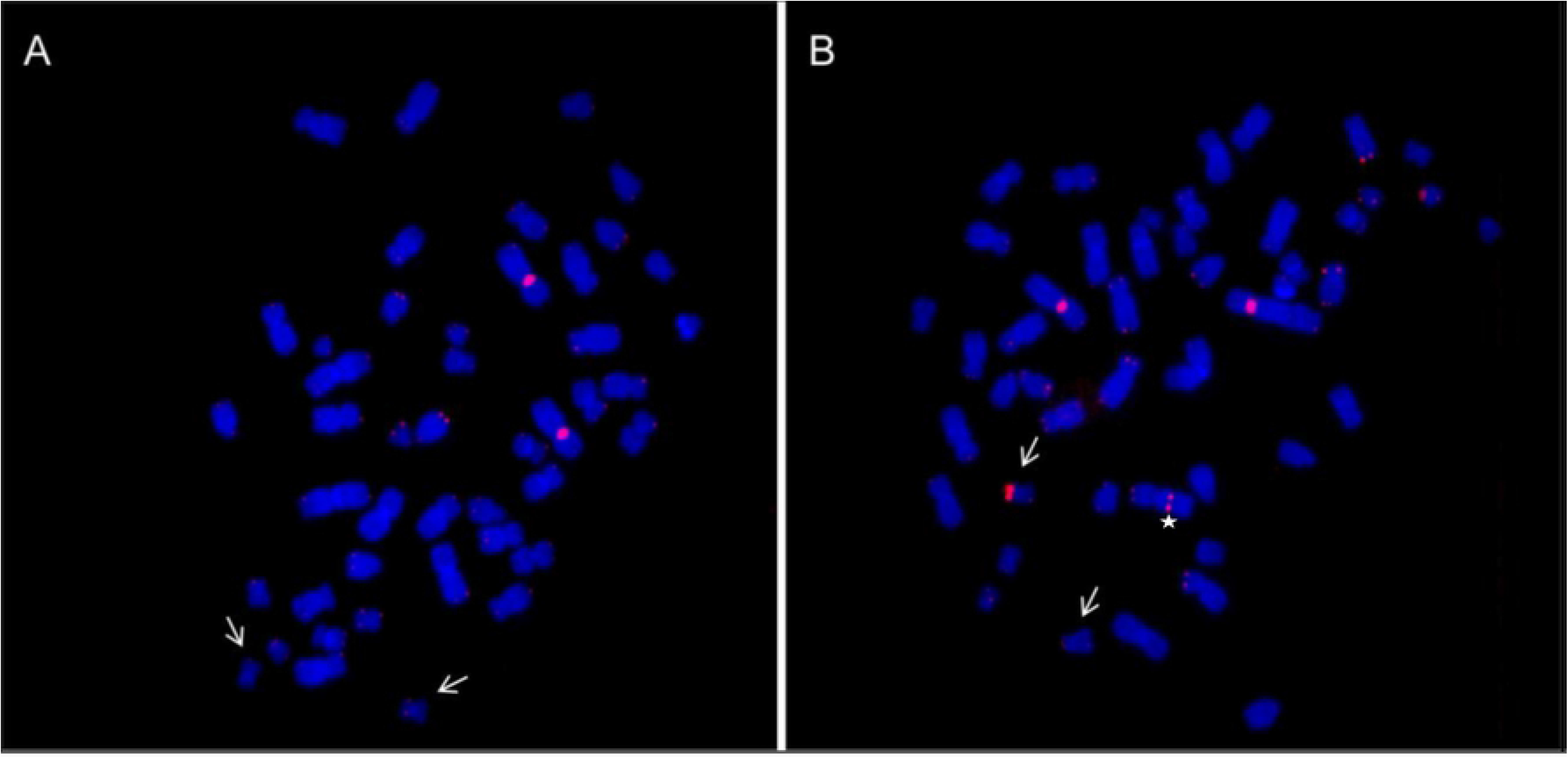
Metaphases of the NBS LCL 94P0307 after Q-FISH. Q-FISH of the NBS cell line 94P0307 with very short telomeres and telomere fusion (×). The arrows point to chromosomes 19. A: Metaphase with weak telomere fluorescence of both chromosomes 19; B: Metaphase with one chromosome 19 with a brightly fluorescent telomere of the p-arm. The bright signal is the reference region of chromosome 2. Original from Habib, 2012[31].

Based on supplemental Fig. 3, the average T/C values can be transformed into qPCR values. Under the assumption that Q-FISH values of 4.8 and 56.6 roughly correspond to qPCR values of 0.1 and 2.5, the absolute TRF lengths are about 11.3 and 18.6 kb, and the length difference about 7 kb. A highly positive correlation was found between telomere lengths estimated by Q-FISH and by qPCR with a correlation coefficient of r = 0.96 (Supplemental Fig. 3). The comparison between the qPCR data and the absolute telomere length measured by telomeric restriction fragments (TRF) analysis resulted in r = 0.64 (Supplemental Fig. 4). In contrast to qPCR, TRF analysis also includes the subtelomeric sequences. To transform the qPCR data into absolute length a regression analysis was applied. The formula is y= 3.4 x X +10.11, where y is the predicted TRF value in kb, X the qPCR value, 3.4 the slope and 10.11 the y-intercept, a constant. In this case, to each qPCR an equivalent in kb of absolute length can be assigned, e.g. if the T/S value according to qPCR is 1.0, the absolute length, y, is 3.4 x 1 + 10.11=13.51 kb.

### Biochemical characterization of the lymphoblastoid NBS cells

The frequency of malignancy may increase in individuals with short telomeres – due to genetic instability - and in individuals with long telomeres in presence of high telomerase activity and immortality. It is thought that that telomere length in normal individuals remains centered around a “malignancy valley,” which on the whole minimizes the rate of cancer [24]. We therefore measured hTERT mRNA levels in our NBS cells. The expression of the telomerase reverse transcriptase gene, h*TERT*, responsible for telomere elongation and induction of immortality, was weak in all lymphoblastoid lines, and was close to the lower detection limit (Supplemental Figs. 1,2).

Caspase 7, a member of effector caspases (caspase 3, 6, 7), is involved in apoptosis and inflammation. Apoptosis plays an important role in protecting organisms from cancer development by eliminating DNA-damaged cells. On the other hand, senescence induction triggered by telomere attrition, is a frequent response to many anticancer modalities and may compensate for attenuated apoptosis. We were therefore interested whether or not differences between the NBS patients cell lines are detectable.

We used activation of caspase as an indicator of the apoptosis capacity. This is indicated by the formation of the Caspase-7 fragment, measured up to 48h after bleomycin treatment. Caspase activity was significantly higher in the three cell lines derived from patients with shorter survival compared to those with longer survival and shorter telomeres (Table 4). The cell line 94P0307 with the shortest telomeres and the high number of telomere fusions displayed an extremely low (almost undetectable) caspase activity.

**Table 4:** Analysis of Caspase-7 after bleomycin treatment of six NBS lymphoblastoid cell lines. Each experiment was repeated three times: Median values plus standard deviation.

|  | Caspase-7 activity after Bleomycin treatment<br>for 0 to 48h |  |  |  |
| --- | --- | --- | --- | --- |
|  | 0h | 12h | 24h | 48h |
| 89P0319 ♀ | 0.65±0.26 | 0.77±0.22 | 0.78±0.17 | 0.90±0.13 |
| 96P0616 ♀ | 0.17±0.01 | 0.24±0.01 | 0.41±0.05 | 0.90±0.08 |
| 95P0182 ♀ | 0.17±0.14 | 0.25±0.15 | 0.46±0.11 | 0.65±0.10 |
| 97P0614 ♂ | 0.13±0.03 | 0.16±0.05 | 0.36±0.09 | 0.49±0.05 |
| RoZd ♀ | 0.13±0.08 | 0.15±0.01 | 0.27±0.01 | 0.36±0.05 |
| 94P0307 ♂ | 0.01±0.01 | 0.02±0.01 | 0.02±0.01 | 0.03±0.02 |

**Table 5:**
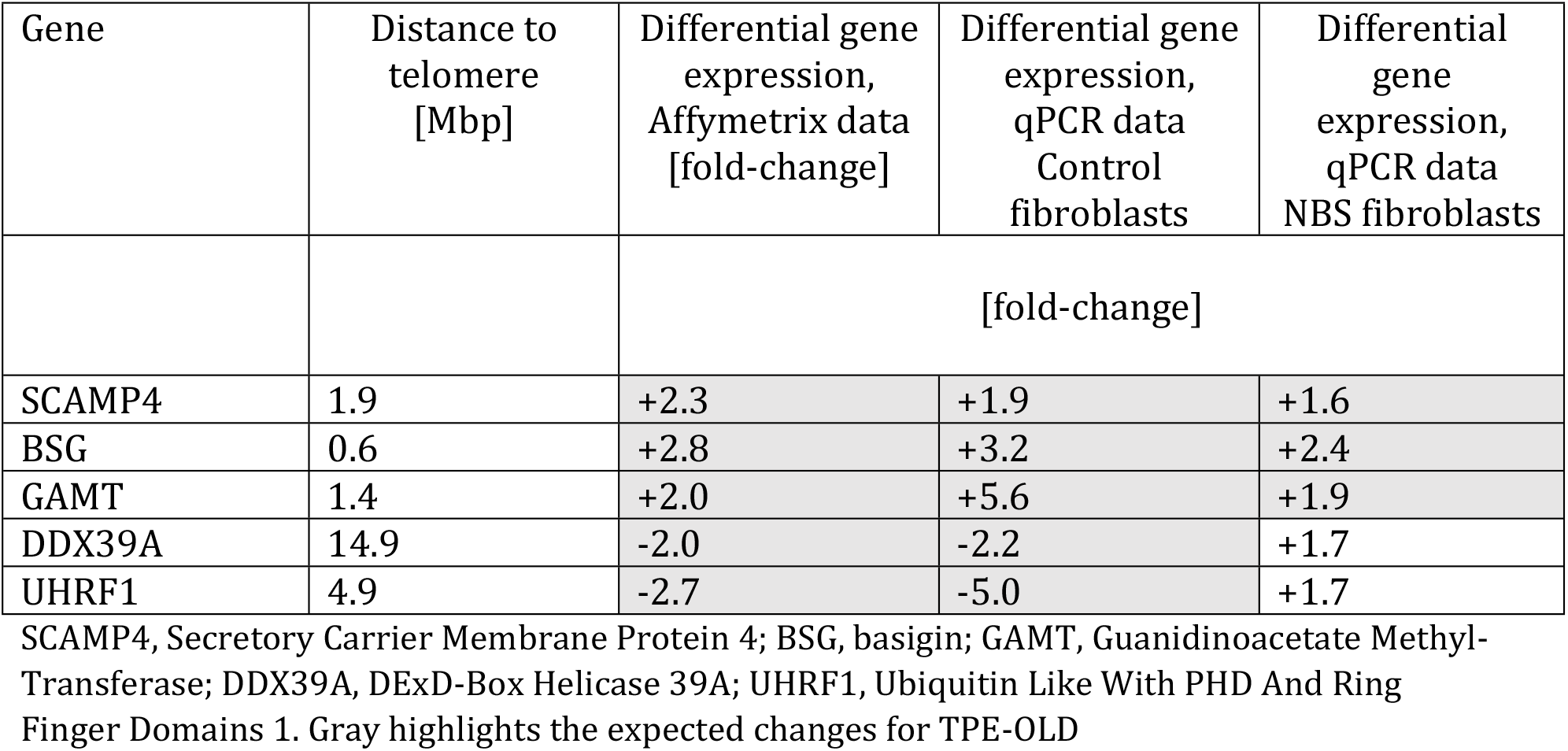
Induction/Suppression of telomeric 19p genes in normal and NBS fibroblasts with short telomeres (relative to cells with long telomeres in hTERT immortalized cells)

| Gene | Distance to telomere [Mbp] | Differential gene expression, Affymetrix data [fold-change] | Differential gene expression, qPCR data Control fibroblasts | Differential gene expression, qPCR data NBS fibroblasts |
| --- | --- | --- | --- | --- |
|  |  | [fold-change] |  |  |
| SCAMP4 | 1.9 | +2.3 | +1.9 | +1.6 |
| BSG | 0.6 | +2.8 | +3.2 | +2.4 |
| GAMT | 1.4 | +2.0 | +5.6 | +1.9 |
| DDX39A | 14.9 | -2.0 | -2.2 | +1.7 |
| UHRF1 | 4.9 | -2.7 | -5.0 | +1.7 |

SCAMP4, Secretory Carrier Membrane Protein 4; BSG, basigin; GAMT, Guanidinoacetate Methyl-Transferase; DDX39A, DExD-Box Helicase 39A; UHRF1, Ubiquitin Like With PHD And Ring Finger Domains 1. Gray highlights the expected changes for TPE-OLD

There is a close functional link between ataxia-telangiectasia and Nijmegen breakage syndrome gene products, ATM and nibrin [36]. Phosphorylation of nibrin, induced by ionizing radiation, requires catalytically active ATM. We therefore also measured ATM phosphorylation at serine-1981. ATM function, however, was not significantly different among cell lines with short or long survival rates. Also the cell line with low caspase 7 activity had normal ATM function, estimated by ATM phosphorylation 1h after bleomycin treatment (Supplemental Table 6).

### Possible implications of ALT and TPE-OLD process in cell line 94P0307

Recently, it has been demonstrated that telomeres may loop to specific loci to regulate gene expression, a process termed TPE-OLD (telomere position effect—over long distance) [37]. In the examples characterized so far, genes close to the telomeres are silenced in young cells (with long telomeres) and become expressed when telomeres are short. Re-elongation of short telomeres in cells by exogenous expression of the h*TERT* gene (active telomerase) results in expression patterns similar to those in young cells with long telomeres. The effect may likely extend to a distance of at least 10 Mbp (upto 15 Mbp) from the telomere and is clearly distinct from classic TPE, which regulates genes proportional to the proximity to the telomeric repeats and is not effective over distances more than 100 Kb from the telomeres [38]

We were therefore interested whether or not the long 19p telomere of the NBS lymphoblastoid cell line 94P0307 might influence the mRNA expression of telomeric genes. For this purpose, we investigated the following TPE-OLD gene candidates of 19p: DDX39A, SCAMP4, BSG (specifically upregulated in pre-senescent cells), and UHRF1 and GAMT (specifically downregulated in senescent cells). In Affymetrix cDNA microarray analyses using normal and Hutchinson-Gilford Progeria fibroblasts, these genes were induced or repressed on the mRNA level in replicative senescence (short telomeres) relative to control cells (with long telomeres), but were not influenced in stress-dependent senescence induced by UV-B radiation (unpublished data). The genes code for proteins involved in epigenetic regulation (SCAMP4), extracellular matrix metalloproteinase induction (BSG), activation of fatty acid oxidation (GAMT), alteration of RNA structure (DDX39A) and cell cycle dependent gene expression (UHRF1).

We investigated relative mRNA levels of these genes with qPCR, using one unrelated fibroblast cell line without and with artificially elongated telomeres in the presence of hTERT, one NBS fibroblast cell line without and with artificially elongated telomeres in the presence of hTERT, and all 6 lymphoblastoid cell lines. As shown in Figure 8 and Table 4, differential regulation in array experiments was confirmed with qPCR for all 5 genes in unrelated healthy control cell lines. In the NBS fibroblast cell line, DDX39A and UHRF1 did not show the expected down regulation at the mRNA level, comparing cells without and with hTERT. The lymphoblastoid cell line with the long 19p telomere did not show any targeted variation relative to the other lymphoblastoid cell lines without extreme telomere elongation. We also found no correlation with the clinical phenotype, comparing three lymphoblastoid cell lines derived from individuals with extremely short survival after cancer manifestation (96P616, 95P182, 59P319), and three from individuals with long survival (RoZD, 97P614, 94P307). Altogether, these data did not support the suggestion that the very long 19p telomere may have influenced TPE-OLD candidate genes in this patient under the experimental conditions described here.

**Fig. 8:**
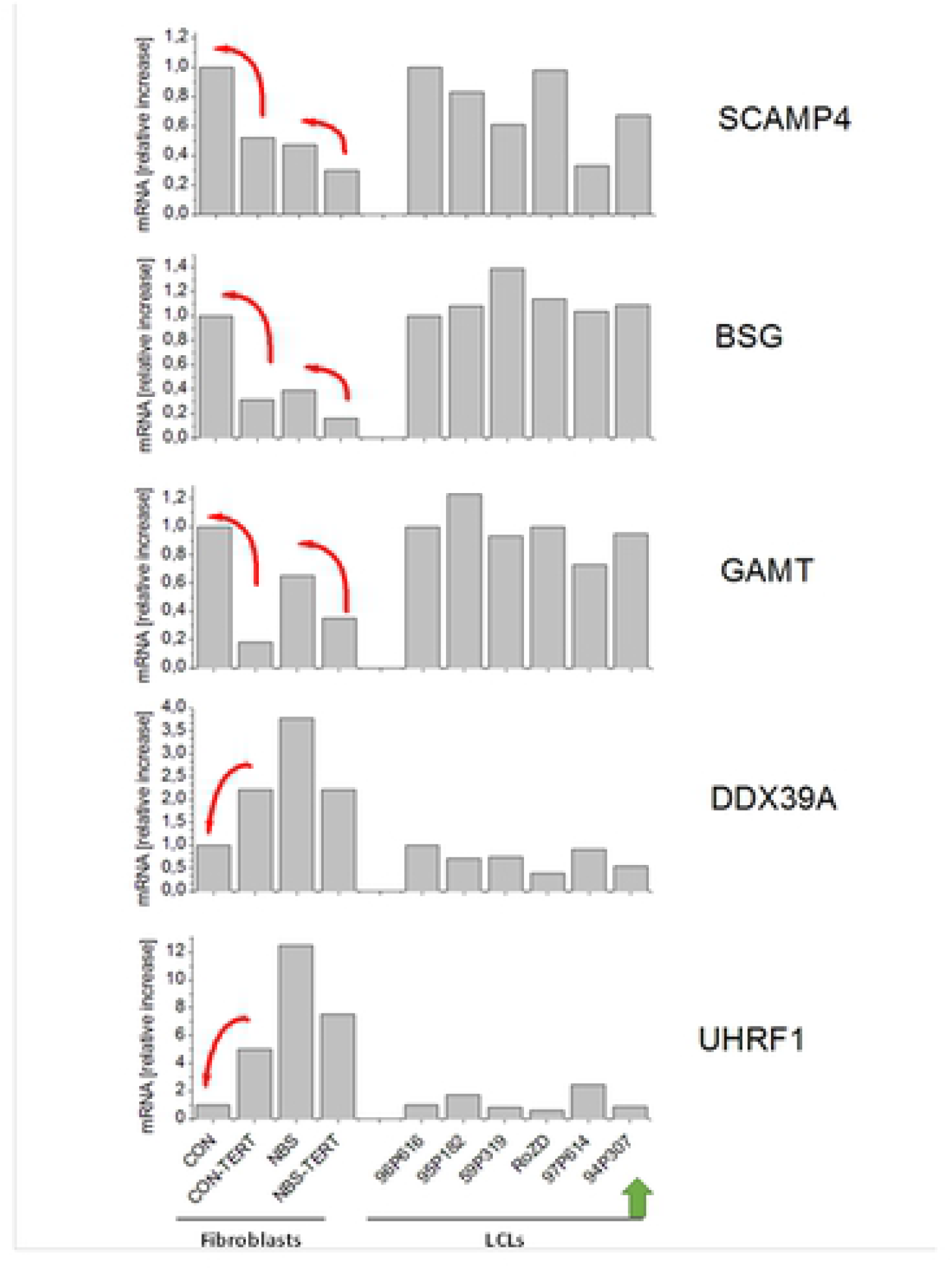
Relative telomere length of three tissues of humanized Nbs mice estimated by qPCR. Relative telomere length (T/S ratio) in brain, heart, and liver tissues of humanized Nbs mice (*Nbn^-/-^NBN^del^*:Nbn^-^): 87, 68 and 92 (age: 32, 41, and 39 days) and humanized wild-type mice (*Nbn^-/-^NBN^+^* : Nbn^+^): 95 and 98 (age: 32 days) estimated by qPCR.

### Telomere length in three tissues of humanized Nbs mice as analysed by qPCR

The common human NBS1 mutation, used in this investigation, was hypothesized to be hypomorphic, so that some low-level functionally relevant truncated nibrin protein may be formed due to an alternative initiation of translation. In agreement with this possibility, null mutation of the homologous gene, Nbn, is lethal in mice. However, Nbn-/- mice with the human NBS allele recapitulate most of the human NBS phenotypes, albeit at a less severe manifestation, including the cell-cycle checkpoint defects [32]. The humanized mice analyzed in this investigation express the human *NBN* gene, generated by the introduction of the human allele with the 5 bp deletion into Nbs-deficient mice (*Nbn*^-/-^*NBN*^del5^). Two humanized Nbs mice with the human wild-type allele (*Nbn*^-/-^*NBN*^+)^ served as control. The T/C ratio of one of the control mice (no. 98) was used as a reference and set to “1”.

Relative telomere length in all tissues of this control mouse was lower than in mice with the mutated allele, but higher in comparison to the other control (no. 95; Fig. 9). Altogether, there was no evidence for a significant difference in telomere length between the mice with the NBN founder mutation and mice with the wild type allele. Thus, longer basal telomere length in mice may have impact on the phenotype and may have prevented a marked telomere attrition in this NBS model.

**Fig. 9:**
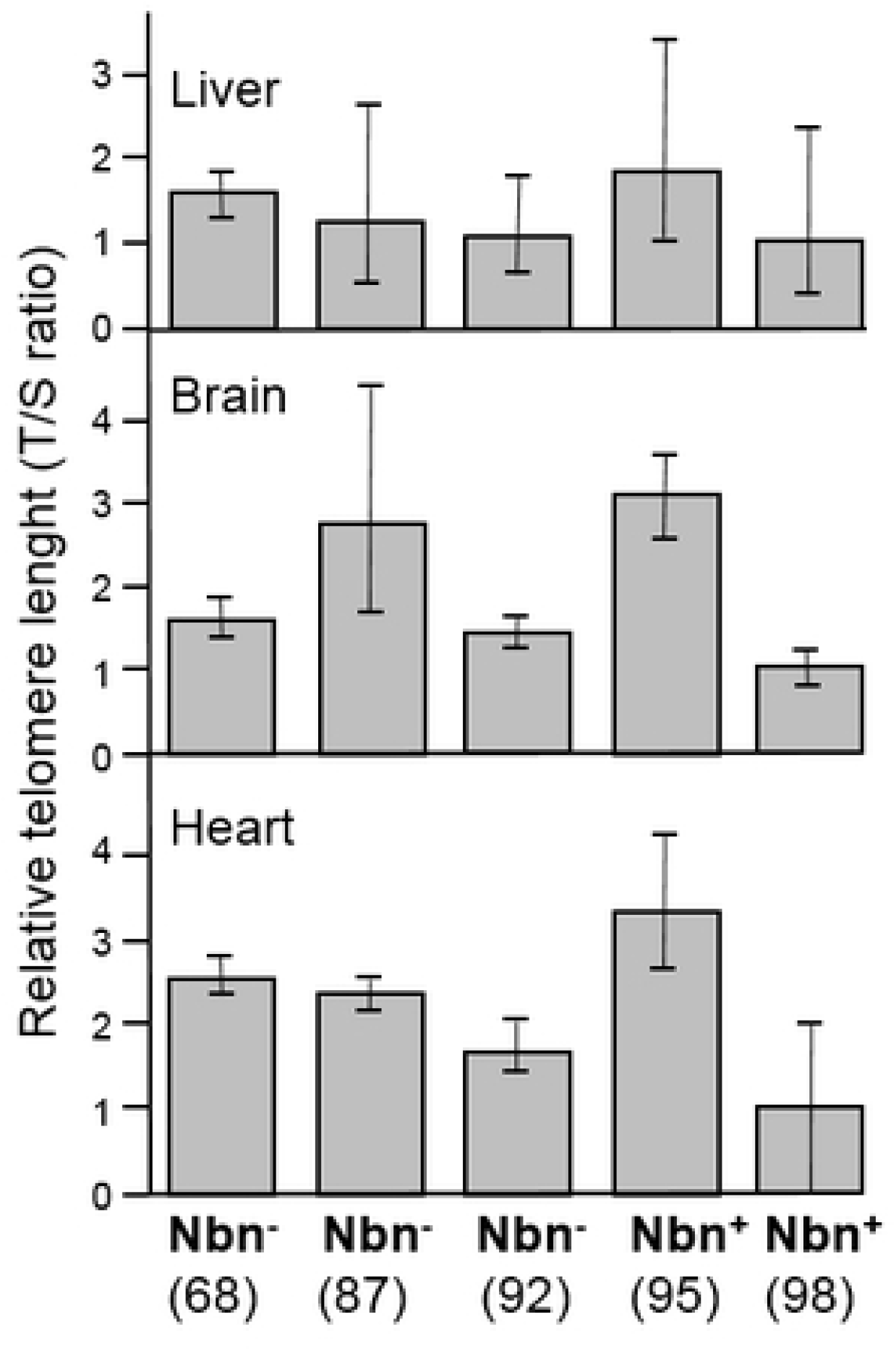
qPCR analysis was performed in a healthy human fibroblast cell line, in a NBS fibroblast cell line, and in all 6 available lymphoblastoid cell lines (LCLs), as described in Table XX. Total mRNA was extracted from proliferating fibroblasts and from the same cell lines proliferating with experimentally elongated telomeres, after immortalization with the catalytic subunit of human telomerase (hTERT). In lymphoblastoid cell lines mRNA from the 6 different donors was compared. One of these cell lines (94P307) showed extremely elongated telomeres on p19. The genes investigated in this experiment were identified as TPE-OLD candidates by use of Affymetrix gene chip experiments in independent cell lines from healthy controls and Hutchinson-Gilford Progeria patients (as described in Table 2). All values were normalized to the level (=1-fold) of mRNA in unmodified and pre-senescent control fibroblasts. Each assay was performed in triplicate. Thin arrows mark the expected trend for TPE-OLD, the thick arrow marks the LCL cell line with extremely long telomeres. SCAMP4, Secretory Carrier Membrane Protein 4; BSG, basigin; GAMT, guanidinoacetate methyltransferase; DDX39A, DExD-Box Helicase 39A; UHRF1, Ubiquitin Like With PHD And Ring Finger Domains 1.

## Discussion

For the first time, this study has systematically examined telomere lengths and related abnormalities in a representative cohort of NBS patients. One characteristic structural DNA change of most dividing cells, *in vivo* and *in vitro*, is the shortening of telomeres, consistent with the “end replication problem” of telomeres and likely acting as tumor-suppressor mechanism in large long-lived species. In addition to considerable individual variability of telomere length in peripheral blood cells, telomere shortening occurs at different rates throughout life. Shortening is most rapid from birth to 4 years of age, slows down between 4 and 39 years and remains at a relatively constant rate through the rest of adult life [39-41]. In our control cohort, the age group of 1-10 years had the longest telomeres with a mean of 16.19 kb, declining to 14.53 kb in the 11-20 yearś age group, remaining rather constant from 21-49 years with a mean of 13.62 kb followed by a decline between 50 and 80 years with a mean of 11.91 kb (calculated by the formula on p. 9). No significant difference in telomere length was found between the control individuals and NBS-heterozygotes younger than 30 years (P = 0.2). In the age group older than 30 years, telomere length was significantly shorter in NBS heterozygotes than in the controls (P = 0.01), though their average age with 42 years was lower than that of the controls with 57.6 years. This points to a more rapid decline in telomere length with age in NBS heterozygotes, paralleled by their significantly increased cancer risk at higher age [9].

NBS homozygotes showed significantly shorter telomere length in all age groups (P<0.05), when compared to controls. This is in line with previous reports in individual cases [19-22] and the findings in other chromosomal instability syndromes [24,25]. We have recently shown that defective nibrin in NBS leads to high levels of reactive oxygen after DNA damage and that this increased oxidative stress is caused by depletion of NAD^+^ due to hyperactivation of the strand-break sensor, Poly(ADP-ribose) polymerase. We therefore hypothesized that the extremely high incidence of malignancy among NBS patients is the result of the combination of a primary DSB repair deficiency with oxidative DNA damage [15]. The results presented here complement this concept in that increased telomere shortening can further contribute to chromosomal instability and increase tumor formation. The telomere shortening and damage is likely the result of disturbed nibrin/MRE11/RAD50 telomere maintenance complex; the increased oxidative damage described by us in the earlier study [15] also potentially damages the telomere structure so that both factors synergistically may increase the cancer risk.

There was, however, no correlation between telomere length and cancer manifestation in the patient group. By contrast, the three patients with the shortest telomeres in lymphoblastoid cell lines showed the longest survival times (>12 years) after cancer manifestation. On the one hand this finding was surprising and contrasts to the concept that telomere attrition may predispose to chronic disease and (at extreme shortening) to genetic instability [42]. On the other hand, the limit on cellular proliferation in cells with short telomeres is considered an initial block to oncogenesis [24]. Extremely short telomeres may prevent an even worse phenotype when a tumor has developed. Remarkably, the cell line with the shortest overall telomere length had an extremely low caspase 7 activity, suggesting that (relative) senescence-associated tumor suppressor effects might be associated with rather low apoptosis rates. Low hTERT expression and the uniform telomere attrition may also have contributed to a relatively stable cellular state and long survival after cancer onset in these individual patients.

NBS is characterized by persistent DNA damage both intrachromosomally and at the telomeres, which results in a pool of cells with deranged genomes prone to cancer manifestation. Not only the average telomere length, but also the chromosomal distribution of telomere shortening may influence the cellular and clinical phenotype. Chromosome-specific features may control telomere length [43-45]. In addition, allelic variation of telomere length has been described [40, 43], as well as subpopulations with different telomere lengths [46]. Using Q-Fish, differences in telomere length were observed between individual chromosome arms, both in NBS and control lymphoblastoid cells. Rather short telomeres were found on chromosomes 17, 19, and 20. However, this pattern is also a characteristic of normal diploid cells [30, 47-49] and no preferred shortening of a particular chromosome was observed in this study.

In the NBS lymphoblastoid cell line 94P0307, which had the shortest total telomere length and a telomere fusion, the telomere of the p arm of one chromosome 19 was brightly fluorescent in 70 % of the metaphases after Q-FISH. The ratio of the T/C values between the chromosomes 19 with strong and weak fluorescence intensity was approximately 11:1. To the best of our knowledge such an allelic variation of a single human telomere has not been previously reported. This observation was remarkable insofar as ALT requires the activity of the MRE11/RAD50/NBS1 recombination complex [50] and NBS1 has been shown to be required for the production of extrachromosomal telomeric circles in human ALT [51] and ALT-associated leukemia bodies [52]. On the other hand, ALT is complex and may also occur without forming T-loops by processes involving homologous recombination and telomere-sister chromatid exchange [50]. The effect of a special ALT mechanism is further strengthened by the fact that we did not find a hint for suppressive chromatin loops or disengagement of suppressive effects in NBS lymphoblastoid cells with short telomeres. Cells with such long telomeres at the end of chromosome 19p are expected to form a chromatin loop in the region of regularly important gene loci [37]. In accordance with this assumption, we could find expected changes on mRNA level in control fibroblasts for all investigated TPE-OLD candidate genes on 19p. We did not, however, find such mRNA changes for the lymphoblastoid cell line despite extremely long telomeres. Altogether these data show that ALT may occur in NBS and that nibrin is not necessarily required for ALT. This observation is in accordance with clinical findings showing cancers with high ALT prevalence in NBS such as sarcoma and glioma [53].

The reduced telomere length in NBS homozygotes was already visible before birth and was in the NBS fetus apparently not related to environmental factors. There were, however, considerable differences in telomere lengths between different fetal tissues: spinal cord and brain tissues had the longest telomeres, fibroblasts and skin the shortest. This is in contrast to the observation of Youngren et al. [54] and Butler et al. [55] who found similar telomere lengths in most fetal tissues. Obviously, the elevated rate of telomere attrition and the tissue differences in NBS fetal telomere length cannot be explained by the “end replication problem” alone. Deficient DNA repair of lesions induced by oxidative stress at telomeres together with S-phase-specific exonucleases, exacerbating the 3’-overhang length [25], may explain an elevated rate of telomere attrition in fetal tissues. Tissue-dependent differences have also been suggested for other telomere disorders (although tissue examinations in these syndromes are scarce) and may explain generational phenotype differences [24]. In accordance with an important role of the MRN complex during development, complete disruptions of NBS1, RAD50 or MRE11 in mice lead to embryonic lethality [32]. Nonetheless, the relatively long telomeres in brain cells is rather unexpected in view of the fetal microcephaly. Microcephaly may be secondary to increased apoptosis [56], which may explain preferable elimination of cells with short telomeres in NBS nerve cells.

Expression of the main human NBS allele rescues the lethality of Nbn-/- mice, and leads to immunodeficiency, cancer predisposition and cell-cycle checkpoint defects associated with a partial reduction in Atm activity [32]. As shown here, these mice displayed some variability in telomere length but did not show overall accelerated telomere attrition. Telomere biology has intensively been studied in the mouse and differs significantly from humans [57]. The telomeres of laboratory mice are about 10 times longer and knock-out of the telomerase gene has almost no effect in the following generation [58]. This effect in combination with the individual variation in telomere length might explain why we observed no reduction in telomere length in the Nbs mice. This fits well with the observation that the disease phenotype in the Nbs mice is less severe than in NBS patients [32] and suggests that telomere attrition may further aggravate the pathology in human beings. On the other hand, these mice have some clinical phenotype in absence of marked telomere attrition, indicating that telomere attrition is rather a secondary event in NBS.

We did not study Nbs mice with artificially shortened telomeres here. Our study has also some other limitations. The TPE-OLD candidate approach does not exclude subtle effects of the long 19p telomere on gene expression that were not detectable under the experimental conditions described here. Cells with the long 19p telomere might have been overgrown by other cells due to a selective disadvantage during cultivation. Moreover, we identified all TPE-OLD candidates in human fibroblasts, and telomeric regulation may be different in lymphoblastoid cells. Interestingly, even two telomeric genes in NBS fibroblasts were not regulated as predicted, which could indicate general dysfunction in inducing telomeric genes in NBS. However, this was not the focus of these investigations and further studies are necessary to investigate the possible role of telomere dysfunction on gene regulation and clinical phenotype in NBS. Finally, we originated our lymphoblastoid cell lines via EBV-infection, which may have influenced telomere function. On the other hand, EBV infects the majority of world population and any effect on telomeres may also have impact for NBS. EBV has the ability to transform B-cells [59,60] and NBS patients develop mainly lymphoid malignancies [61].

In summary, our data show that telomere length is dramatically reduced in multiple cells and tissues in NBS, and that telomere attrition is likely associated with alterations in telomere function and elongation. According to recent recommendations for a novel nomenclature [24], NBS can best be described as a secondary telomeropathy. Our data do not currently warrant the classification of the disease as a classical “impaired telomere maintenance spectrum disorder” (ITM). Secondary telomeropathies may include other diseases with defects in the DNA repair machinery, such as FANCD2 and RECQL4, as well as mutations in genes with poorly characterized telomere function, such as MPN1 and DNMT3B. In these disorders it is not clear if the telomere dysfunction is the cause or just one of the effects of the diseases. Enhanced telomere attrition further contribute to chromosomal instability and high cancer risk in NBS. The long survival times of individual NBS patients with extreme telomere abrasion most likely reflect the bifunctional role of telomere shortening. Moreover, our data suggest that nibrin is not necessarily required for ALT. The exact mechanism and clinical significance of ALT in absence of nibrin as well as implications for tumor progression have to be evaluated further.

## Material and Methods

### Cell cultures

The cell lines were established with the ethical approval of “The Children’s Memorial Health Institute, Warsaw” and the Ethics Committee of the 2^nd^ Medical School of the Charles University, Prague. The fibroblast cell lines were propagated in Amniomax medium with gentamicin sulfate and L-glutamine supplement. Lymphoblastoid cells (LCLs) were prepared as previously described [27] and cultured in RPMI medium with L-glutamine (RPMI+glutamax™) supplemented with 15% FBS (<u>F</u>etal <u>B</u>ovine <u>S</u>erum), and the usual amount of antibiotics. Cells were grown in incubators at 37°C and 5% CO_2_.

### DNA samples

DNA was isolated at the Institute of Medical and Human Genetics, Berlin from blood of 27 NBS heterozygotes, 38 NBS homozygotes, from a homozygous fetus, and from 108 controls.

The fetus with NBS presented with microcephaly as detected by ultrasound. Amniotic cells showed a 46, XY karyotype, and molecular diagnostics revealed homozygosity for the NBS founder mutation. The pregnancy was interrupted in the 32^nd^ week of pregnancy. The fetal weight was below normal for gestational age (25^th^ percentile), with craniofacial dysmorphology, signs of immaturity such as short nails and missing *“*Béclard’s nucleus”, but no malformations of inner organs. The head circumference was below the 3^rd^ percentile, the brain was slightly hypotrophic without malformations. After obtaining informed consent, tissues were obtained for further histological and genetic investigations.

### Molecular genetics

The founder mutation was analysed by PCR of exon 6 of the *NBN* gene and sequencing, as previously described [6] Expression of the human telomerase reverse transcriptase gene (hTERT) was analysed by qPCR. The hTERT cDNA was synthesized according to the standard manufacture’s instruction (Invitrogen). GAPDH (glyceraldehyde-3-phosphatedehydrogenase) was used as internal control gene. All PCRs were performed on Applied Biosystem prism 7500 (software DSD V1.2.3). The qPCR products were checked by agarose gel electrophoresis, visualized by UV-transilluminator, and photographed.

For mRNA quantification of TPE-OLD (telomere position effect over long distances) candidate genes, the total RNA was isolated using the extraction kit from Macherey-Nagel (NucleoSpin RNA, Cat. 740955) and quantified in a NanoDrop ND 1000 spectophotometer. cDNA synthesis was carried out following the manufacturer’s instructions (M-MLV Reverse Transcriptase, RNase (H-), Promega, Cat: M5301). 1 µg mRNA was used for the reverse transcriptase reaction. qPCR was carried out using TaqMan Universal Master Mix II, no UNG (Cat. 4440040) from Applied Biosystemes following the manufacturer’s manual. The cDNA samples (10 ng/µl) were assayed in triplicates; non template control (water) was analyzed in duplicates in a 384 – well plate. The assay was run on a BioRad CFX384 real – time C1000 thermal cycler with the following thermal cycling profile: 10 min at 95°C, followed by 50 cycles at 95°C for 15 s and 1 min at 60°C with signal acquisition. For gene expression quantification the following gene expression TaqMan assays from Applied Biosystems were used: SCAMP4 (Cat. Hs00365263_m1), BSG (Cat. Hs00936295_m1), GAMT (Cat. Hs00355745_g1), DDX39A (Cat. Hs01124952_g1), UHRF1 (Cat. Hs01086727_m1). Human PPIA (Cyclophilin A) (Cat. 4333763F) was used as endogenous control. The ΔΔCt method was used for relative quantification, as previously described [28]. The relative expression level of each gene was normalized to the proliferating pre-senescent control fibroblasts.

Quantitative polymerase chain reaction (qPCR) was applied to determine the relative telomere length as described by Cawthon (2002) [29]. The *36B4* gene, located on chromosome 12, was chosen as reference single copy gene. The number of telomere copies (T) in relation to the single copy gene (S) reflects the relative telomere length (T/S ratio). As a reference for all qPCR measurements a mix of blood DNA from three healthy individuals (two females, one male) of the same age (32 years) was used and set as “1”.

Quantitative fluorescence in situ hybridization (Q-FISH) of telomere repeats was performed according to Perner et al. (2003)[30] to determine the relative length of individual telomeres. To do so, the fluorescence intensity *of single* telomeres (T) relative to a constant repetitive region in the centromeric region (C) of chromosome 2 (T/C ratio) was measured. In addition, the total relative telomere length of all chromosomes was estimated. In short: Metaphases are captured and karyotyped using a florescence microscope with individual filter sets (DAPI and Cy3), and linked to Isis and Ikaros software from Metasystems. The telomere measurement algorithm displays two overlaid horizontal lines on each chromosome in the karyogram. These define the telomere measurement areas for the p- and q-arms. The area of the reference probe is displayed in the same way (Fig. 1). The T/C ratio of 15 metaphases is estimated for each case. The mean of telomere intensities of the p- and q-arms, and standard deviation, were calculated for each chromosome.

Absolute telomere length was measured by Terminal Restriction Fragment (TRF) length analysis, performed according to the manufacturer’s instructions (Roche - TeloTAGGG telomere length assay). The protocol involved DNA fragmentation using a combination of the frequently cutting restriction enzymes Hinf1 and Rsa1. Fragments were subsequently resolved by gel electrophoresis, and transferred to a nylon membrane. The blotted DNA fragments were hybridized to a digoxigenin (DIG)-labeled probe specific for telomeric repeats, incubated with a DIG-specific antibody, exposed to an x-ray film to estimate the mean TRF length. Detailed protocols of the above cytogenetic and molecular genetic methods, apart from the TPE-OLD approach, are presented in Habib, 2012 [31].

Humanized NBS mice, kindly provided by A. Nussenzweig, were generated by reconstitution of Nbn knockout mice with the human *NBN* gene carrying the c.657_661del5 mutation or the wild type allele[32]. Three different tissues (heart, brain, and liver) were analysed from three humanized NBS mice (*Nbn*^-/-^*NBN*^del5^) at the age of 32, 39, and 41 days and two control individuals (*Nbn*^-/-^*NBN*^+^) at the age of 32 days. The mouse studies were approved by the *Landesamt für Gesundheit und Soziales*, Berlin (G0438/09).

### hTERT immortalization

Fibroblasts were infected with retroviral supernatants from a packaging cell line (PA317-TERT), which stably expresses the human telomerase cloned into a pBabePuro vector, according to reference. The vector and packaging cell line were a friendly gift from Dr. Woodring Wright (UTSW Medical School, Dallas, USA). After a two-week selection phase with puromycin, the infection and selection were controlled by determination of hTERT expression using real-time PCR and measurement of telomere length [33,34]

### Identification of TPE-OLD candidate genes in human fibroblasts

Using Affymetrix cDNA microarray we compared the expression profiles of two different types of aging models: fibroblasts from patients suffering from Hutchinson-Gilford progeria (HGP) and fibroblasts, in which senescence was induced by UV radiation (using four 20W TL/12 lamps emitting broadband UV-B peaking at 312 nm.; twice a day for 3 consecutive days with a total dose of 528 J/m²). The unnormalized raw data of all experiments were exported in an ASCII format from the Affymetrix GCOS®-Software and further analysed by the programme iReport from Ingenuity. We analyzed 16 microarrays, each with 54.675 transcripts on an HGU133-A2.0 array from Affymetrix®, in a design with 3 groups (HGP vs. controls; UB-B-treated controls vs. controls; HGP vs. HGP-TERT). The analysis was based on pairwise comparison algorithm, standard two-sided *t*-tests and *F*-tests to find differentially expressed genes (*α* < 0.05). Differentially expressed genes (DEGs) in HGP, upto 15 Mbp telomeric on p19, that were NOT DEG in UV-B treated controls <u>and</u> were NOT DEG in HGP-TERT were viewed as potential TPE-OLD candidates (Supplementary Figure). Detailed whole-genome analysis will be published elsewhere.

### Analysis of Caspase-7 in Western Blots

Lmyphoblastoid cells in logarithmic growth were treated in triplicate with 10 μg/ml Bleomycin for 0h, 12h, 24h, and 48h. After centrifugation the cell pellet was resuspended, stained with Ponceas S and photographed. After destaining the blot was incubated with the primary antibody *cleaved Caspase-7*(Asp 198) against Caspase-7 and the second antibody (Anti-Mouse IgG Horsradish Peroxidase). Exposure and documentation was performed as above. Actin served as internal control.

### Detection of ATM and phosphorylated ATM (p-ATM) by immunoprecipitation

For examination of phosphorylation of ATM after bleomycin treatment lymyphoblastoid cells in logarithmic growth were treated in triplicate with 0, 10 and 30 μg/ml Bleomycin for 1 h at 37°C. The proteins were extracted according to standard procedures. Immunoprecipitation was performed with the FISH-antibody (Anti-ATM, rabbit polyclonal, Novus) and magnetbeads (Dynal Biotech ASA/Invitrogen). The proteins were separated by gel-elektrophoresis, transferred to a nitrocellulose membrane, stained with Ponceas S, and photographed. Thereafter, the membrane was destained by H_2_Odd, probed first with a primary anti-ATM pS1981 antibody (Rochland, monoclonal, mouse), reprobed with an anti-ATM antibody (Abcam Cambrige, UK), and finally with an anti-actin antibody as a loading control (Abcam, Cambridge, UK). Chemiluminescence detection was performed after incubation with the secondary antibody ECL (Anti-Mouse IgG Horsradish Peroxidase linked whole antibody: GE Healthcare UK limited) for 2h at room temperature. After treatment with solution 1(Enhanced Luminol Reagent; Perkin Elmer) and 2 (Oxidizing Reagent; Perkin Elmer) for 1 min, an X-ray film (Kodak Medical X-ray Film, General Purpose Blue) was applied and exposed for different times. Detailed protocols of the caspase and ATM experiments are presented in reference [35].

### Statistical tests

The original data were exported to Excel 2007, GraphPad Prism 5 software and SPSS15.0 software for graphs and boxplots. Statistical tests were Mann-Whitney U test, Fisher’s exact test, and the unpaired T test.

### Ethics statement

All procedures were in accordance with the ethical standards of the responsible committee. The cell lines were established with the ethical approval of “The Children’s Memorial Health Institute, Warsaw” and the Ethics Committee of the 2nd Medical School of the Charles University, Prague and the consent of the proband’s parents. The mouse studies were approved by the Landesamt für Gesundheit und Soziales, Berlin (G0438/09).

## Acknowledgments

R. Habib received a stipend from the Damascus University, Syria; R. Kim received a stipend from the Gottlieb Daimler-und Karl Benz-Stiftung. This work was supported by grants from the DFG to K.S. and M.D. (Collaborative Research Centre 577, Project B1); We acknowledge the probands and their parents for their participation, Annelore Junge for prenatal cytogenetic diagnostics, and André Nussenzweig for providing the humanized Nbs mouse. We thank Antje Gerlach, Mohsen Karbasiyan, Janina Radszewski, and Bastian Salewsky for technical help, Britta Teubner and Brigitte Schröder for her cytogenetic assistance and Dr. Rami Derbas for statistical advice.

## Author contributions

RH analysed the telomere lengths in the framework of her dissertation (Q-PCR, Q-FISH, hTERT analysis). RK performed the caspase and ATM experiments in the framework of his dissertation. KC and ES took care of the probands and performed the primary molecular diagnostics. RF provided fetal data and tissues.. KJ and MW performed the TPE-OLD (telomere position effect over long distances) analysis and hTERT immortalization; ID the TRF analysis. HN established and characterized the lymphoblastoid cell lines; MD provided resources; KS and MW drafted the paper and all authors read and accepted the manuscript.

## Ethics approval and consent to participate

All procedures were in accordance with the ethical standards of the responsible committee. The cell lines were established with the ethical approval of “The Children’s Memorial Health Institute, Warsaw” and the Ethics Committee of the 2^nd^ Medical School of the Charles University, Prague and the consent of the proband’s parents. The mouse studies were approved by the *Landesamt für Gesundheit und Soziales*, Berlin (G0438/09).

## Author details

Dr. Raneem Habib Department of Human Genetics, Ruhr-University Bochum, Bochum, Germany

Dr. Ryong Kim Institute of Clinical Chemistry, Red-Cross General Hospital, Pyongyang, Democratic People’s Republic of Korea

Prof. Dr. Heidemarie Neitzel Institute of Medical and Human Genetics, Charité - Universitaetsmedizin Berlin; Berlin, Germany;

Prof. Dr. Ilja Demuth Lipid Clinic at the Interdisciplinary Metabolism Center, Charité - Universitaetsmedizin Berlin, Berlin, Germany

Prof. Dr. Krystyna Chrzanowska Department of Medical Genetics, The Children’s Memorial Health Institute, Warsaw, Poland

Prof. Dr. Eva Seemanova Department of Clinical Genetics, 2^nd^ Medical School, Charles University, Prague, Czech Republic

Prof. Dr. Renaldo Faber Zentrum für Pränatale Medizin Leipzig, Germany

Prof. Dr. Martin Digweed Institute of Medical and Human Genetics, Charité - Universitaetsmedizin Berlin; Berlin, Germany;

Dr. Kathrin Jäger Institute of Clinical Chemistry and Laboratory Medicine, University of Rostock

Prof. em. Dr. Karl Sperling Institute of Medical and Human Genetics, Charité - Universitaetsmedizin Berlin; Berlin, Germany;

Prof Dr. Michael Walter Institute of Clinical Chemistry and Laboratory Medicine, University of Rostock

## Supplemental Information

**Supplemental Table X1:** Telomere length of healthy individuals (controls)

**Supplemental Table X2:** Estimation of telomere length of NBS-homozygotes

**Supplemental Table X3:** Estimation of telomere length of NBS-heterozygotes

**Supplemental Table X4:** Correlation between telomere length and clinical data of 21 NBS homozygotes.

**Supplemental Table X5:** T/C-FISH data on telomere lengths of NBS-LCL 94P0307

**Supplemental Table X6:** ATM phosphorylation after 1h Bleomycin treatment in six NBS lymphoblastoid cell lines

**Supplemental Fig. 1:** Expression of the hTERT gene in lymphoblastoid cells. Agarose gel electroporesis of the PCR products of the hTERT cDNA and the GAPDH cDNA as internal control. 06P0131 and 96P0125 are lymphoblastoid controls derived from male individuals homozygous for the wild type allele. Original from Habib 2012[31].

**Supplemental Fig. 2A:** Expression of hTERT as measured by qPCR. Dissociation curve of qPCR for *hTERT* cDNA (template): Original from Habib 2012[31]

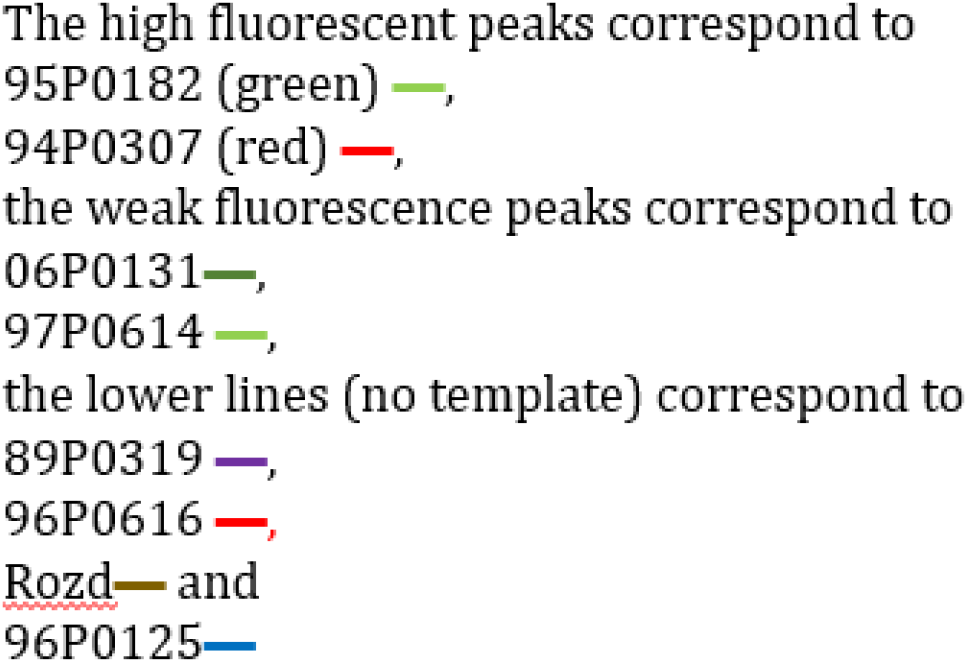

Original from Habib 2012[31]

**Supplemental Fig. 2B:** Expression of hTERT as measured by qPCR in a separate experiment. Dissociation curve of qPCR for *hTERT* cDNA (template) and non template control (NTC):

**Supplemental Fig. 3:** Correlation between telomere length analysed byqPCR and Q- FISH in six NBS lymphoid cell lines and two controls (06P0131,96P0125). Original from Habib 2012[31].

**Supplemental Fig. 4:** Correlation of telomere length measured by qPCR and TRF analysis. The equation of the computed regression line, which is used to correlate the data, is y= 3.4 x X +10.11 with a correlation coefficient of r= 0.64.

